# CaMK2rep: A Highly Sensitive Genetically Encoded Biosensor for Monitoring CaMKII Activity in Mammalian Cells

**DOI:** 10.1101/2025.05.07.652623

**Authors:** Elena Martínez-Blanco, Raquel de Andrés, Lucía Baratas-Álvarez, F. Javier Díez-Guerra

## Abstract

Accurately monitoring calcium/calmodulin-dependent protein kinase II (CaMKII) activity in cells remains a significant challenge due to the limited sensitivity and narrow dynamic range of existing genetically encoded sensors. Here, we introduce CaMK2rep, a novel phosphorylation-based biosensor that enables robust, specific, and high-sensitivity detection of CaMKII activity. CaMK2rep is designed with two tandem CaMKII consensus sites embedded within the native sequence context of synapsin, and its phosphorylation is detected via a phospho-specific antibody, allowing both biochemical and morphological analyses. We validated CaMK2rep in HeLa cells and cultured hippocampal neurons, demonstrating a near-linear response to CaMKII expression levels and stimulation intensity, and no detectable cytotoxicity. To complement CaMK2rep measurements, we employed the live-cell CaMKAR^1^ reporter to monitor CaMKII activity dynamics. Using both tools, we investigated the role of neurogranin (Ng), a major postsynaptic calmodulin (CaM) binding protein, and obtained consistent evidence supporting a CaM-buffering model in which Ng limits basal CaMKII activation by sequestering CaM. These findings establish CaMK2rep as a sensitive, specific, and versatile biosensor for CaMKII signaling, particularly well-suited for immunoblot-based population analyses. They also illustrate the value of combining orthogonal genetically encoded tools to interrogate complex signaling mechanisms in both physiological and pathological contexts.

## Introduction

Calcium/calmodulin-dependent protein kinase II (CaMKII) is a multifunctional serine/threonine kinase that phosphorylates a wide range of substrates and plays a central role in intracellular Ca²⁺ signaling, particularly in neurons and cardiac tissue^2,3^. The CaMKII family comprises four isoforms –α, β, γ, and δ-each encoded by distinct genes, with alternative splicing further expanding their functional diversity^4^. In the brain, CaMKIIα and CaMKIIβ are predominant, with CaMKIIα expressed at approximately threefold higher levels^5,6^. CaMKIIα is the most abundant protein at excitatory synapses, accounting for up to 10% of the total protein in the postsynaptic density (PSD)^7–9^. It is essential for synaptic plasticity mechanisms such as long-term potentiation (LTP) and dendritic spine remodeling, both of which underlie learning and memory^10–13^. CaMKIIα monomers assemble into 12-subunit homo-or heteromeric holoenzymes, with each subunit retaining kinase activity^14^.

Historically, CaMKIIα was viewed as a molecular memory switch, due to its autophosphorylation at Thr286 (pThr286) which confers autonomous activity that extends beyond transient Ca^2+^ signals. Recent evidence challenges the idea that pThr286 is necessary for the long-term memory maintenance^15^. Instead, pThr286 appears critical for the induction of various plasticity forms –including LTP, long-term depression (LTD), and behavioral timescale synaptic plasticity (BTSP)-but dispensable for their long-term persistence. The current view proposes that a calcium transient initiates T286 autophosphorylation and triggers the translocation of CaMKII to the postsynaptic density (PSD), where it interacts with GluN2B^16^ and may also fulfill a structural role^17^.

Given the central role of CaMKII in intracellular signaling, particularly in neurons, there is a need for reliable tools to monitor and to manipulate its activity with high specificity Current approaches include pharmacological inhibitors and genetically encoded reporters^18^. Among the latter, Camuiα –a FRET-based sensor-has been widely used to study CaMKII dynamics in live cells^19,20^. Camuiα is composed of a CaMKIIα backbone flanked by cyan (CFP) and yellow (YFP) fluorescent proteins, enabling detection of conformational changes associated with Ca^2+^/calmodulin (CaM) binding and autophosphorylation at Thr286 (pThr286). Upon activation, Camuiα exhibits a decrease in FRET efficiency. Despite its great utility, Camuiα presents several limitations. It operates with low FRET efficiency^21^ and requires overexpression of a modified CaMKIIα, which may not faithfully reflect endogenous enzyme activity or dynamics. Its integration into native holoenzymes is uncertain and could be impaired by the bulky fluorescent tags. Moreover, Camuiα primarily senses conformational shifts rather than directly reporting on kinase activity, making it difficult to distinguish between Ca^2+^/CaM binding and pThr286-driven autonomous activity. It is also unresponsive to ATP-competitive inhibitors and may fail to capture pThr286 dephosphorylation events following LTP induction^22^. Finally, Camuiα is restricted to the α isoform and does not report on the activity of other CaMKII variants.

To address these limitations, the FRET-based sensor FRESCA^23^ was developed to monitor endogenous CaMKII activity without altering its expression levels. FRESCA uses a CaMKII-specific substrate (syntide-2) and a phosphothreonine-binding domain (FHA2) to detect phosphorylation events. Unlike Camuiα, FRESCA does not incorporate CaMKIIα itself, avoiding interference with native holoenzyme composition. However, it exhibits a much lower dynamic range, with signal changes approximately ten times smaller than those of Camuiα. Although both FRESCA and Camuiα can theoretically be targeted to specific subcellular compartments, the addition of localization sequences at either end of the polypeptides may alter FRET efficiency and would require further validation. More recently, a novel CaMKIIα activity reporter named CaMKAR was introduced^1^. CaMKAR employs a circularly permuted GFP (cpGFP) fused to the CaMKII autoregulatory peptide (MHRQE**T**VDCLK).. Its phosphorylation induces a conformational change that increases GFP fluorescence, enabling ratiometric detection. Unlike Camuiα, CaMKAR responds to both catalytic and allosteric CaMKII inhibition, providing broader functional coverage. In HEK293T cells, it outperforms Camuiα with a tenfold higher signal-to-noise ratio and a threefold faster response. Nevertheless, its dynamic range remains modest, reaching a maximum 2.5-fold increase in fluorescence upon activation.

The present study introduces CaMK2rep, a novel substrate-based reporter, specifically designed to improve the sensitivity and specificity of CaMKII activity detection. CaMK2rep consists of a nuclear export signal (NES), a green fluorescent protein (sGFP2), a tandem repeat of the rat synapsin-1a phospho-site type 3 (S603), and three myc tags. CaMK2rep demonstrated to be highly sensitive and selective for CaMKII. While not yet applicable for live-cell imaging, it effectively complements existing CaMKII biosensors by offering enhanced sensitivity in fixed samples or endpoint assays. We used CaMK2rep to investigate the regulatory role of Neurogranin (Ng), a CaM-sequestering protein, in modulating CaMKII activity. Our results show that Ng attenuates CaMKII activity following Ca²⁺ influx in both HeLa cells and cultured hippocampal neurons. Moreover, studies using Ng mutants which do not bind CaM binding confirmed that Ng’s effect on CaMKII activity depends on its ability to sequester CaM. These findings were validated using the CaMKAR sensor. We discuss these results in light of previous studies proposing a regulatory role for Ng in shaping synaptic excitability. In summary, CaMK2rep is a new sensitive and specific reporter of CaMKII activity that broadens the current toolkit for investigating CaMKII function, especially useful in high-throughput assays. This study advances our understanding of the mechanisms underlying synaptic homeostasis by highlighting the modulatory role of Ng in synaptic plasticity.

## Materials and Methods

### Animals and ethics compliance

Wistar rats were bred at the animal facility of the Universidad Autónoma de Madrid (UAM). All procedures conducted during the study strictly adhered to the Spanish Royal Decree 1201/2005, which governs the protection of animals used in scientific research, as well as the European Union Directive 2010/63/EU concerning the welfare of animals in scientific contexts. Additionally, all experimental protocols received approval from both institutional and regional ethics committees, ensuring that the highest standards of animal care and ethical compliance were maintained throughout the research.

### Reagents

Fetal bovine serum (FBS), Dulbecco’s modified Eagle’s medium (DMEM), 0.25% trypsin, Neurobasal media and B27 supplement were from Thermofisher Scientific. The protease inhibitor cocktail was from Biotools (B14001). Total protein was measured using the Bradford Protein Assay kit (Bio-Rad). Pre-stained protein markers VI (10-245 kDa) were from PanReac-AppliChem. Immobilon-P membranes and ECL western blotting reagent were from Millipore. Oligonucleotides and synthetic DNAs were purchased from Integrated DNA Technologies (IDT). 1-beta-arabino-furanosylcytosine (AraC) was from Calbiochem (251010). N-methyl-D-aspartic acid (NMDA), UBP684 and UBP714 (NMDAR PAMs) and Bicuculline methiodide were from Hello Bio. Paraformaldehyde (PFA) was from Merck. PEI Max transfection reagent (MW 40,000) was from Polysciences (Cat. n° 24765). Antibodies and plasmids used are listed in Supplementary Tables 1-3.

### CaMK2rep construct design and cloning

To generate the various CaMK2rep constructs, we synthesized a 499 bp DNA fragment (SynP3myc; IDT) containing the following elements (Supp. Fig. 1A): (i) 43 bp corresponding to the C-terminal region of mCherry; (ii) a 237 bp sequence encoding amino acids 543–620 of rat synapsin-1a, in which the original CaMKII-specific phospho-site 2 (QATRQASISGP, residues 560–570) was replaced with phospho-site 3 (PIRQASQAGPG, residues 598–608), resulting in two tandem phospho-site 3 motifs within this segment; and (iii) three tandem myc tags (EQKLISEEDL). The SynP3myc fragment was PCR-amplified and cloned into the pRSET-mCherry vector using Gibson assembly (NEB). A second synthetic fragment (Spot-NES, 303 bp; Supp. Fig. 1A) encoding a Spot-tag (PDRVRAVSHWSS), a nuclear export signal (NES: SRLQLPPLERL), and 135 bp of the mCherry C-terminus was inserted at the N-terminus of mCherry in the pRSET-mCherry-SynP3myc vector via Gibson assembly. The complete construct (Spot-NES-mCherry-SynP3myc) was then subcloned into the pcDNA3.1 vector and mCherry replaced by superGFP2^24^ (sGFP2), yielding pcDNA3.1-Spot-NES-sGFP2-SynP3myc. This plasmid was renamed CaMK2rep and used for transfections in HeLa cells.

For expression in primary neurons, the NES-sGFP2-SynP3myc cassette was cloned into a pLOX lentiviral (LV) vector^25^ under the control of the human synapsin promoter. To enhance postsynaptic targeting, the PSD95.FingR cDNA (309 bp; Addgene #119736), which encodes an intrabody that binds endogenous PSD-95^26^, was inserted upstream of NES-sGFP2-SynP3myc sequence in the LV vector by Gibson assembly. PSD95.FingR markedly improved the peripheral localization of the reporter in cultured hippocampal neurons (Figure 4B). Four lentiviral constructs were generated (with/oPSD95.FingR and with/o sGFP2) and evaluated for expression and phosphorylation efficiency in primary neurons. Among these, the construct termed nCaMK2rep3 (35 kDa) was selected for its robust expression, strong phosphorylation, and unique ability to co-immunoprecipitate with PSD-95 (Figure 4C). This construct was renamed nCaMK2rep to distinguish it from CaMK2rep, the pcDNA3.1-based reporter. Site-directed mutagenesis (Ser to Ala) was used to generate versions of both CaMK2rep and nCaMK2rep containing only one or no CaMKII phosphorylation sites, which served as negative or partial phosphorylation controls.

### Culture and transfection of HeLa cells

HeLa cells were maintained in Dulbecco’s Modified Eagle’s Medium (DMEM) supplemented with 10% fetal bovine serum (FBS), without antibiotics. Cells were passaged at 90% confluence using 0.25% (w/v) trypsin, 0.53 mM EDTA solution, twice weekly, at a split ratio of 1:3 to 1:6. For transfection, cells were seeded at a density of 15,000 cells/cm² to reach 70–80% confluence on the day of transfection. Transient transfection was carried out using PEI-MAX. Cells were incubated for 5 hours at 37 °C with 5% CO₂ in OptiMEM (Gibco) containing PEI MAX at a ratio of 2 µL PEI-MAX per 1 µg DNA. After incubation, the transfection medium was replaced with fresh 10% FBS/DMEM. Assays were typically conducted 24-hour post-transfection. All plasmids were purified using the Qiagen Midiprep kit.

### Primary cultures of rat hippocampal neurons

Primary hippocampal neurons were prepared from embryonic day 19 (E19) rat embryos as previously described^27^. Embryos were collected and maintained in chilled incomplete Hank’s balanced salt solution (HBSS) during dissection. After brain removal, hippocampi were carefully isolated and freed of meninges. Tissue was washed five times in incomplete HBSS, then incubated with 0.25% trypsin for 15 minutes at 37 °C to facilitate dissociation. After trypsin removal by two additional washes in incomplete HBSS, the tissue was transferred to complete HBSS containing DNase I (0.04 mg/mL) and dissociated mechanically using Pasteur pipettes and 22G needles. The resulting cell suspension was filtered through a 70 µm nylon mesh, centrifuged at 1200 rpm for 5 minutes, resuspended in plating medium, and counted. Cells were plated at a density of 25,000 cells/cm² onto culture dishes pre-coated with 0.1 mg/mL poly-L-lysine (PLL) in borate buffer (pH 8.0), or at 12,000 cells/cm² on coverslips pre-treated with 0.25 mg/mL PLL. Coverslips were previously sterilized with 65% nitric acid (24–72 h), extensively washed with double-distilled H_2_O (ddH_2_O) and baked at 180 °C. 3 hours after seeding to allow adhesion to the substrate, the plating medium was replaced with Neurobasal medium supplemented with B27 and GlutaMAX (Thermofisher). To suppress glial proliferation, 1 µM AraC was added on day in vitro 3 (DIV3). On DIV7, 50% of the culture medium was replaced with fresh Neurobasal + B27. Cultures were maintained at 37 °C in a humidified atmosphere with 5% CO₂ and used for experiments during the third week in vitro. All treatments were performed within the thermostatized CO_2_ incubator.

### Preparation of lentiviral and adeno-associated viral particles

#### Lentiviral particles (LVs) production

HEK-293T cells (3 × 10^6^) were seeded in p100 dishes and transfected 24 hours later with 8 µg of the lentiviral plasmid of interest, 4 µg of pCMVδR8.74, and 2 µg of pMD2.G using PEI MAX (DNA:PEI ratio 1:2). DNA and PEI were mixed in OptiMEM medium (Gibco), incubated for 20 minutes at room temperature (RT), and added dropwise to the cells. After 5 hours of incubation at 37 °C and 5% CO₂, the transfection medium was replaced with Neurobasal medium. Viral supernatants were collected 48 hours post-transfection, filtered through a 0.45 µm filter, aliquoted, and stored at –80 °C. Hippocampal neurons were infected at day in vitro 4 (DIV4).

#### Adeno-associated virus (AAV) production

AAVs were prepared as described by McClure et al. 2011^28^. HEK-293T cells (7 × 10⁶) were seeded in p150 dishes and cultured in 10% FBS/DMEM until 70% confluency, at which point the medium was replaced with 5% FBS/DMEM. Cells were then transfected using the calcium phosphate method with the following plasmids: 12.5 µg of the desired pAAV plasmid, 25 µg of pFδ6, 6.25 µg of pH21, and 6.25 µg of pRV1. Plasmids were diluted and adjusted to a volume of 900 µL using CaCl_2_ (final concentration 125 µM). An equal volume of 2× HBS was added, the solution bubbled and incubated for 20 minutes at RT, then added dropwise to the cultures. After 16 hours of incubation (37 °C, 5% CO₂), the medium was replaced by 10% FBS/DMEM. 48 hours post-transfection, cells were washed with PBS and lysed in extraction buffer (150 mM NaCl, 0.4% sodium deoxycholate, 50 U/mL benzonuclease (Millipore), 20 mM Tris-HCl pH 8). Lysates were centrifuged at 3,000 × g for 15 minutes, and the supernatant was applied to a Heparin Sepharose column (HiTrap, Cytiva). Eluted fractions were concentrated using Amicon Ultra-4 100 kDa filters (Millipore), filtered through 0.13 µm filters, aliquoted, and stored at −80 °C. Hippocampal neurons were typically infected at DIV7 with AAV particles diluted in half the culture volume of the dish. After an 8-hour incubation, the viral medium was replaced with a 1:1 mixture of the previously collected medium and fresh Neurobasal medium supplemented with B27 and GlutaMAX. To silence Neurogranin (Ng) expression, the plasmid shNg pA_RC3J1_CAGW (Addgene #92155), encoding an shRNA targeting the sequence gtgacaagacttccctactgt, was used. Prior to shNg AAV production, the GFP coding region of the original plasmid was replaced by mRuby2. As a control, a non-targeting scrambled shRNA (gtgccaagacgggtagtca, Addgene #181875) was used. To preserve neuronal viability, AAV infections for knockdown experiments were performed at DIV10.

### Protein extraction and Western blots

Cells were lysed in extraction buffer containing 50 mM NaCl, 0.5% Triton X-100, 1 mM EDTA, 2 mM DTT, 25 mM Tris-HCl (pH 6.8), and protease & phosphatase inhibitors (Biotool). Lysates were homogenized by 20 passes through a 23G needle and centrifuged at 17,500 × g for 15 minutes at 4 °C. Protein concentrations in the supernatants were determined using the Bradford assay (Bio-Rad). For SDS-PAGE, 5–10 μg of protein from HeLa cell extracts or 15–25 μg from hippocampal neuron extracts were loaded per well. Proteins were separated by SDS-PAGE on 10% or 13% polyacrylamide gels (Mini Vertical Protein Electrophoresis System, Cleaver Scientific) under reducing conditions at 120 V for ∼2.5 hours. Protein in the gels was transferred onto PVDF membranes (Millipore) using a semi-dry blotting system (Nyx Technik) at 400 mA for 30 minutes in transfer buffer (22.5 mM Tris, 170 mM glycine, 20% methanol). Membranes were blocked with 5% (w/v) skimmed milk in TBS for 1 hour at room temperature with agitation and incubated overnight at 8 °C with primary antibodies in TBS with 0.05% Tween-20. HRP-conjugated secondary antibodies (Jackson ImmunoResearch, 1:15,000) were used for detection with an enhanced chemiluminescence system (ECL, Millipore). Signal acquisition was performed using an Amersham Imager 680 (GE Healthcare Life Sciences), and densitometric analysis was carried out with the open source software Fiji/ImageJ^29,30^.

### Immunoprecipitation

Primary hippocampal neuron cultures were washed with cold PBS, lysed in extraction buffer, homogenized by 20 passes through a 23G needle, and centrifuged at 17,500 × g for 15 minutes at 4 °C. Protein concentration in the supernatant was determined using the Bradford assay (Bio-Rad). 10% of the lysate was reserved as input, and the remaining was incubated overnight at 8 °C with rotation in the presence of anti-myc antibody (1 μg per mg of protein). Subsequently, 30 μl of a 50% (v/v) Protein G-agarose suspension (ABT) was added and incubated for 1 hour at 4 °C. The beads were then washed three times with cold lysis buffer, resuspended in 40 μl of Laemmli buffer, heated at 95 °C for 5 minutes, and analyzed by Western blot using a mouse anti-PSD95 antibody (Millipore, MAB1596, clone 6G6).

### Immunofluorescence of cultured cells

HeLa cells or primary hippocampal neurons (DIV16–17) grown on 18 mm round coverslips were quickly washed with PBS and fixed with 4% paraformaldehyde (PFA) in PBS for 20 minutes at room temperature. Residual aldehydes were quenched by incubation with 0.2 M glycine (pH 8) in PBS for 5 minutes. After three PBS washes, cells were permeabilized and blocked for 30 minutes in blocking buffer containing PBS, 0.1% Triton X-100, 1% bovine serum albumin (BSA), and 1% heat-inactivated horse serum. Primary antibody incubation (Supplementary Table 3) was carried out overnight at 8 °C in PBS supplemented with 1% BSA and 1% horse serum. After three 5-minute PBS washes, secondary antibodies were applied for 1 hour at room temperature in the same buffer. Nuclei were counterstained with DAPI (0.2 μg/mL in PBS) for 5 minutes, followed by washes in distilled water and 96% ethanol. Coverslips were air-dried and mounted using Mowiol. Images were acquired on a Zeiss Axiovert 200M fluorescence microscope and processed using Fiji/ImageJ^29,30^. Raw 16-bit monochrome images were denoised using the Noise2Noise plugin^31^, background-subtracted, and color-coded with appropriate lookup tables.

### Photoactivation assay

For HeLa cells, we used the plasmids CMV-tdTomato-P2A-paCaMKIIα (Addgene #165431) and CMV-tdTomato-P2A-paCaMKII(K42M) (Addgene #165433)^32^. For hippocampal neurons, we employed pAAV-CaMP0.4-FHS-paCaMKII-WPRE3 (Addgene #165429) and pAAV-CaMP0.4-FHS-paCaMKII(SD)-WPRE3 (Addgene #165430)^32^. HeLa cells were co-transfected with the CaMK2rep plasmid, while hippocampal neurons were co-infected with nCaMK2rep lentiviral particles. In HeLa cells, the culture medium was replaced with complete Hank’s buffer (135 mM NaCl, 5.3 mM KCl, 1.25 mM CaCl_2_, 0.8 mM MgSO_4_, 5.56 mM glucose, 0.45 mM KH_2_PO_4_, 0.34 mM Na_2_HPO_4_, and 10 mM HEPES, pH 7.4), followed by a 20-minute incubation at room temperature. For hippocampal neurons, 1 μM tetrodotoxin (TTX) was added to the culture medium, and cells were incubated for 30 minutes at 37 °C in 5% CO₂. The medium was then replaced with pre-warmed complete Hank’s buffer supplemented with 1 μM TTX, followed by an additional 20-minute incubation at room temperature. To release paCaMKII activity, cells were continuously illuminated with a 460 nm blue LED (CoolLED pE-4000) at an intensity of 3 mW/cm² for 2 minutes. Cells were then lysed in extraction buffer, homogenized, and centrifuged. Phosphorylation of (n)CaMK2rep in the supernatant was assessed by Western blot.

### Live-cell imaging

HeLa cells were seeded on acid-cleaned, heat-sterilized 25 mm round coverslips and transfected 24 hours later. The next day, coverslips were transferred to Attofluor™ chambers (Thermofisher) and maintained in complete Hank’s medium at 37 °C for 15 minutes prior to imaging. Hippocampal neurons were seeded on poly-L-lysine-coated 25 mm round coverslips at a density of 12,000 cells/cm², and infected at DIV4 and DIV7 with lentiviral (LVs) and adeno-associated viral (AAVs) particles, respectively. At DIV15, the culture medium was replaced with pre-warmed complete Hank’s medium, incubated at 37 °C for 15 minutes, and transferred to Attofluor™ chambers. Unless otherwise noted, imaging was performed using a Zeiss Axiovert 200M fluorescence microscope at 37 °C, equipped with a 25× multi-immersion objective (NA 0.8), a CoolLED pE-4000 light source, and a PCO Edge 4.2 monochrome camera. For calcium imaging, HeLa cells were transfected with pcDNA3-tdTomato-P2A-jGCaMP8s, derived from Addgene #162371. This plasmid encodes the calcium indicator jGCaMP8s^33^ (filters: ex: 470/20 nm; em: 525/50 nm) and the red fluorescent protein tdTomato^34^ (filters: ex: 550/15 nm; em: 580/30 nm). Images were acquired at 4-10 Hz for 5 minutes. jGCaMP8s intensity values were normalized to those of the tdTomato channel, which is insensitive to calcium fluctuations. To monitor CaMKII activity, we used CaMKAR^1^ (Addgene #205315), a genetically encoded CaMKIIα reporter provided by Dr. Jonathan M. Granger (Johns Hopkins University). CaMKAR fluorescence (filter em: 518/35 nm) was recorded with alternating 395 nm (filter ex: 395/25 nm) and 470 nm (filter ex: 470/24 nm) excitation at 1 Hz for 5 minutes. As a negative control, we used CaMKAR-T6A, a phospho-dead mutant that displays no response to CaMKII activation, confirming the specificity of the reporter. CaMKAR responses were quantified as the ratio of 470/395 channel intensities. Image processing and analysis was performed using Fiji/ImageJ^29,30^. Background signals from cell-free regions were subtracted prior to analysis. jGCaMP8s and CaMKAR responses were expressed as ΔF/F_0_, where F_0_ is the mean baseline intensity and ΔF the difference between the fluorescence intensity at each time point and F_0._

### Statistical analysis

Data were analyzed using GraphPad Prism 8. For parametric comparisons, Student’s t-test was used for two groups, and one-way or two-way ANOVA with Bonferroni post hoc correction for multiple groups. For non-parametric comparisons, the Mann–Whitney test or Kruskal–Wallis test was applied for two or more groups, respectively. Results are presented as mean ± SEM from at least three independent experiments. Statistical significance was defined as follows: * p < 0.05, ** p < 0.01, *** p < 0.001, **** p < 0.0001. The absence of asterisks indicates non-significant differences (ns).

## Results

CaMKII, a serine/threonine kinase, is involved in multiple signaling pathways that regulate key physiological processes and contribute to the pathogenesis of neurological and cardiovascular diseases^2^. Its activity is controlled by local Ca^2+^ oscillation amplitude and frequency, the availability of CaM and the dephosphorylation of T286 and T305/306 residues. In the postsynaptic compartment, CaMKII and Ng are highly concentrated relative to CaM. Ng binds CaM in a Ca^2+^– and protein kinase C (PKC)-regulated manner, forming a dynamic complex that shapes CaMKII activation and synaptic plasticity. Two main, non-mutually exclusive models have been proposed to explain Ng’s role in synaptic plasticity: one proposes that Ng facilitates CaMKII activation by recruiting and releasing CaM at synapses upon stimulation^35–37^; the other proposes that Ng, by reducing the binding affinity of Ca^2+^ to the C-terminal lobe of CaM^38^, limits Ca^2+^/CaM-dependent signaling, particularly at low Ca^2+^ levels. To clarify Ng’s functional contribution, we set out to measure CaMKII activity as a readout of synaptic plasticity. Previous attempts using the reporters Camuiα and FRESCA yielded insufficient sensitivity in this context. We therefore developed and validated a new CaMKII activity reporter (CaMK2rep) with enhanced sensitivity to provide deeper insights into Ng’s role in synaptic plasticity.

### Design of CaMK2rep, a CaMKII activity reporter

To design CaMK2rep, we sought CaMKII-specific substrate sequences and focused on Synapsin I, a well-characterized CaMKII target. Among its phosphorylation sites, site 3 (Ser603) is highly specific for CaMKII-mediated phosphorylation^39^ (Figure 1A). We selected amino acids 543–620 of rat Synapsin-1a, which encompass both phospho-site 2 (QATRQASISGP, residues 560–570) and phospho-site 3 (PIRQASQAGPG, residues 598–608). To enhance CaMKII responsiveness, we replaced the sequence of phospho-site 2 with that of site 3, generating a 78-amino-acid polypeptide containing two identical CaMKII-specific phosphorylation motifs. For detection and localization, the polypeptide was fused to superGFP2 (sGFP2^24^) and a nuclear export sequence (NES) at the N-terminus, and three consecutive myc tags at the C-terminus. The full construct was cloned into the pcDNA3.1 vector. To serve as non-phosphorylatable controls, we generated two additional CaMK2rep variants containing Ser603Ala mutations in one or both phosphorylation sites. Phosphorylation of CaMK2rep was assessed by western blot using phospho-specific antibodies: anti-phospho-Synapsin I (pSer603) (Millipore #AB5883) or anti-phospho-Synapsin I (Ser605) (D4B9I, Cell Signaling #88246). The latter provided a stronger signal with lower background and was therefore used in all subsequent experiments. For initial validation, we transfected HeLa cells, which express negligible levels of endogenous CaMKII^40^.

**Figure 1.**
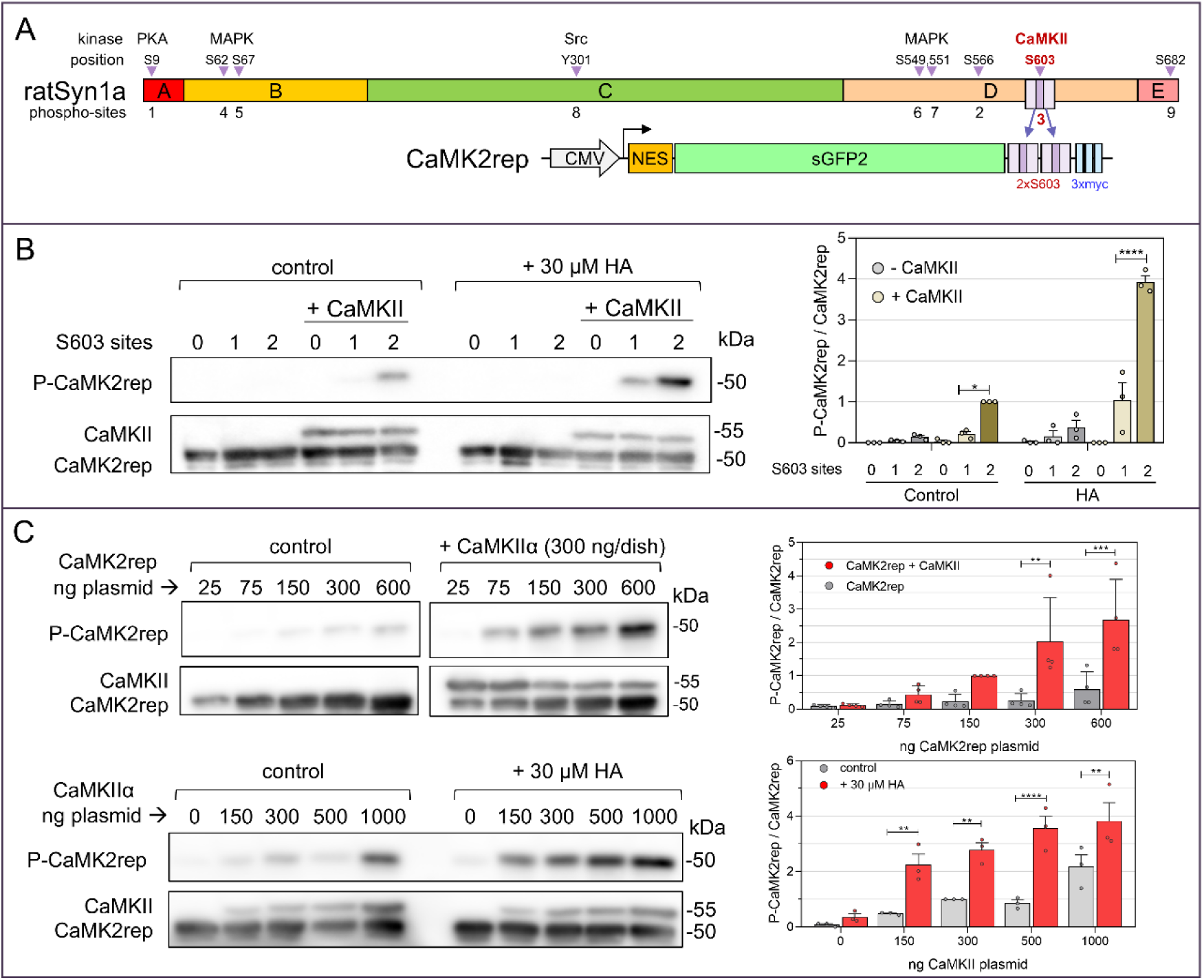
Design and characterization of CaMK2rep in HeLa cells. **A.** Above: Diagram of the domains (A-E) and phosphorylation sites of rat synapsin-1a^39^. Below: Schematic representation of the CaMK2rep, a fusion protein containing a nuclear export signal (NES), super green fluorescent protein 2 (sGFP2), two tandem repeats of the rat synapsin-1a phosphorylation site (S603) and 3 10-aa myc sequences. **B.** HeLa cells transfected with CaMK2rep carrying two, one, or no phosphorylatable Ser603 residues were stimulated with HA (30 µM, 2 minutes) with or without CaMKII expression. Lysates were analyzed by western blot for CaMKII and CaMK2rep levels, and for CaMK2rep phosphorylation. **C.** HeLa cells were transfected with increasing amounts of CaMK2rep with or without CaMKII (300 ng, upper panel) or with 150 ng of CaMK2rep and increasing amounts of CaMKII (lower panel). CaMK2rep phosphorylation was analyzed in non-stimulated cells (upper panel) and after HA stimulation (30 µM, 2 minutes) (lower panel). In all cases, data in the histogram represent the ratios of phospho-CaMK2rep / CaMK2rep and were normalized to the non-stimulated (control) condition with 150 ng CaMK2rep and 300 ng CaMKII (mean ± SEM; B, n=3; C, n=4).

### Characterization of the CaMK2rep in HeLa Cells

When expressed in HeLa cells, CaMK2rep displayed a predominantly cytoplasmic distribution, with no detectable signal in the nucleus or intracellular vesicles (Supplementary Figure 1B). To assess its phosphorylation, we compared CaMK2rep variants containing zero, one, or two Ser603 sites, with or without co-expression of CaMKIIα, under basal conditions and following stimulation with 30 µM histamine (HA) (Figure 1B). In HeLa cells, HA activates histamine H1 receptors, which couple to Gq proteins to stimulate phospholipase C (PLC), leading to inositol triphosphate (IP3) production and calcium release from the endoplasmic reticulum (ER)^41^. Phosphorylation of CaMK2rep was only observed in cells expressing CaMKIIα, even after HA stimulation, confirming the reporter’s specificity for CaMKII activity. As expected, the non-phosphorylatable mutant (zero S603A sites) showed no detectable phosphorylation, underscoring the high specificity of the D4B9I antibody for phospho-site 3. Moreover, the reporter with two Ser603 sites exhibited significantly higher phosphorylation levels than the single-site variant, indicating that duplication of the CaMKII target sequence enhances both the sensitivity and dynamic range of CaMK2rep. To optimize assay conditions, we titrated plasmid concentrations and found that 150 ng of CaMK2rep and 300 ng of CaMKIIα (per 10 cm² dish at 70–80% confluency) provided the best balance between signal strength and dynamic range (Figure 1C). These conditions were used in all subsequent experiments.

We next examined how varying intracellular Ca^2+^ levels affected CaMK2rep phosphorylation by stimulating HeLa cells with different concentrations of histamine (HA) and analyzing the response over time. Intracellular Ca^2+^ levels increased rapidly following HA addition, reaching a peak within the first minute at concentrations of 30 and 100 µM (Figure 2A). As expected, in the absence of CaMKIIα, HA stimulation did not induce detectable phosphorylation of CaMK2rep, further confirming the reporter’s specificity. Among the concentrations tested, 30 µM HA elicited the highest level of CaMK2rep phosphorylation in cells co-expressing CaMKIIα (Figure 2B). Notably, a low level of basal CaMK2rep phosphorylation was observed even without HA stimulation in the presence of CaMKIIα. This likely reflects constitutive or spontaneous CaMKIIα activity under basal conditions, possibly driven by low-level Ca²⁺/CaM signaling. We observed that HeLa cells show small and sporadic calcium transients without external stimulation. Further, previous studies have shown that CaMKIIα can be partially activated when only two of CaM’s four Ca^2+^-binding sites are occupied^42^. We then tested different stimulation periods with HA (30 µM) of 1, 2 or 5 minutes. We found that 1-2 minutes after HA addition were those in which maximal levels of CaMK2rep phosphorylation were obtained (Figure 2C).

**Figure 2.**
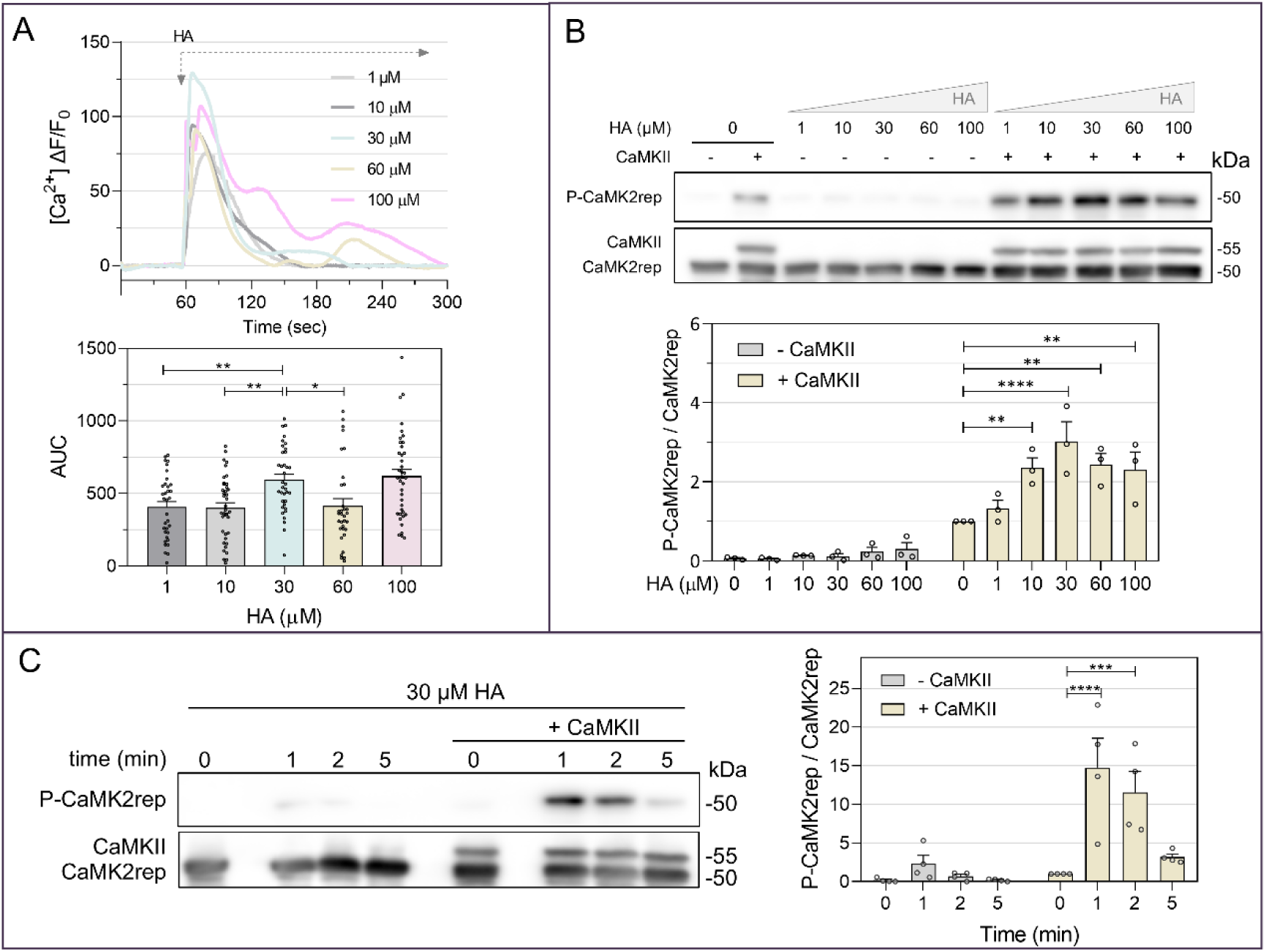
HA increases intracellular calcium levels and CaMK2rep phosphorylation. **A.** Intracellular Ca²⁺ levels measured in HeLa cells in response to HA using the calcium biosensor tdTomato-P2AjGCaMP8s. Images were acquired at 10 Hz using a 25× objective on a Zeiss Axiovert200M fluorescence microscope. The graph shows the mean response calculated from a total of 39–45 cells in three independent experiments. The histogram represents the area under the curve (AUC) for each individual cell computed for 60 sec after HA addition. **B-C.** HeLa cells transfected with CaMK2rep with or without CaMKII were stimulated with increasing concentrations of HA for 2 minutes (**B**), or exposed to 30 μM HA for 1, 2, or 5 minutes (**C**). CaMK2rep phosphorylation levels were normalized to the condition with +CaMKII and no HA (mean ± SEM, n=3 for panel B, n=4 for panel C).

At this stage, CaMK2rep demonstrated high sensitivity to CaMKIIα activity in HeLa cells, with negligible phosphorylation observed in the absence of the kinase, indicating strong specificity. To further validate this specificity, we tested whether other kinases could induce CaMK2rep phosphorylation (Figure 3A). Activation of the cAMP pathway by forskolin, overexpression of the catalytic subunit of PKAα, with or without HA stimulation did not result in detectable CaMK2rep phosphorylation. Similarly, overexpression of PKCγ, in combination with either the PKC activator phorbol-12-myristate-13-acetate (PMA) or the inhibitor calphostin C, failed to elicit any CaMK2rep phosphorylation signal. We next assessed the effect of KN-93, a well-characterized Ca^2+^/CaM-competitive inhibitor of CaMKII^43^, which effectively blocks Ca^2+^/CaM –dependent activation and T286 autophosphorylation but does not interfere with autonomous kinase activity^44^. KN-93 abolished the HA-induced phosphorylation of CaMK2rep in cells co-expressing CaMKIIα, while the inactive analog KN-92 had no effect (Figure 3B), further confirming CaMK2rep’s specificity. To provide additional evidence, we employed photoactivatable CaMKII (paCaMKII), a chimeric protein in which CaMKIIα is fused to the light-sensitive LOV2 domain^32^. In the absence of light, paCaMKII responds to Ca^2+^/CaM like wild-type CaMKIIα, but upon blue light (460 nm) stimulation, it becomes autonomously active, bypassing the need for Ca^2+^/CaM. As shown in Figure 3C, CaMK2rep phosphorylation increased upon light activation of paCaMKII, whereas no phosphorylation was observed with the kinase-dead mutant paCaMKII-K42M. Together, these results provide strong evidence that CaMK2rep selectively reports CaMKII activity with high specificity and minimal cross-reactivity to other kinases.

**Figure 3.**
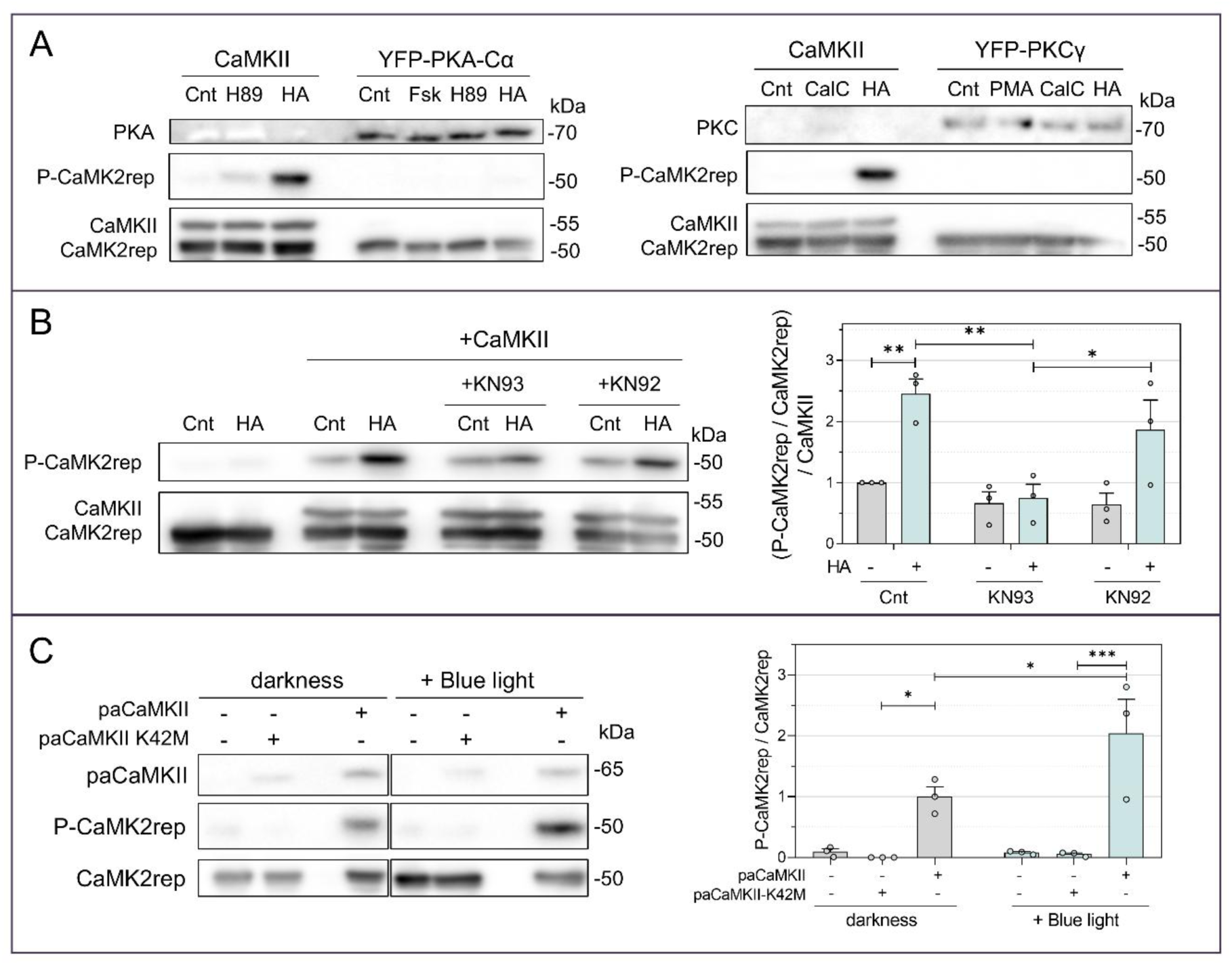
CaMK2rep is specifically phosphorylated by CaMKII activity. **A.** HeLa cells co-transfected with CaMK2rep and the kinases CaMKII, YFP-PKA (left), or YFP-PKC (right), were then treated with one of the following: 30 μM HA 2 minutes, 20 μM Forskolin (Fsk) for 10 minutes, 10 μM H89 for 60 minutes, 100 nM PMA for 10 minutes or 1 μM Calfostin-C (CalC) for 60 minutes. No CaMK2rep phosphorylation was observed in the presence of either PKA or PKC. **B.** HeLa cells co-transfected with CaMK2rep and CaMKII were treated with 20 μM KN-93 or its inactive analog KN-92 for 10 minutes at 37°C, followed by stimulation with 30 μM HA for 2 minutes. CaMK2rep phosphorylation was normalized to the control condition without HA (mean ± SEM, n=3). **C.** CaMK2rep phosphorylation was analyzed after blue light activation of paCaMKII^32^. HeLa cells transfected with CaMK2rep and paCaMKII and stimulated with blue light (460 nm) for 2 minutes showed increased CaMK2rep phosphorylation. The inactive mutant paCaMKII-K42M was used as a control. Mean ± SEM, n=3.

**Figure 4.**
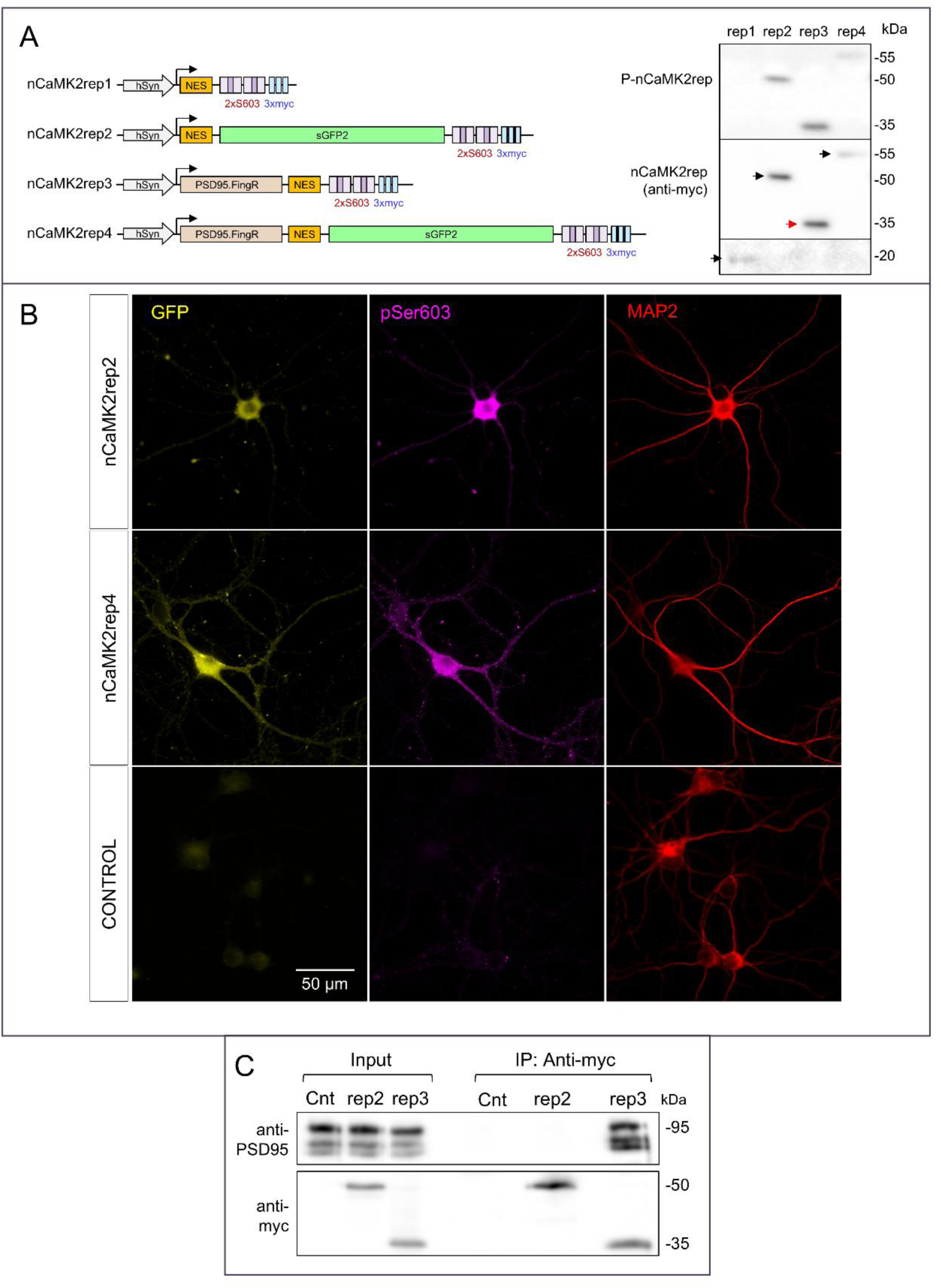
Design of nCaMK2rep for use in primary neuronal cultures. **A.** Schematic representation of CaMK2rep variants in a lentiviral vector (pLOX) under the synapsin promoter. Expression and phosphorylation of each CaMK2rep (nCaMK2rep1-4) were analyzed by western blot of hippocampal neurons infected at DIV4 and lysed at DIV16. **B.** Immunofluorescence of non-infected hippocampal neurons (control) or infected with nCaMK2rep2 and nCaMK2rep4. GFP signal was captured directly in fixed cells with a 63× objective in a Zeiss Axiovert 200M fluorescence microscope. Anti-MAP2 and anti-pS603 (D4B9I) were used as described in Methods. **C.** Lysates of neurons (DIV16) expressing nCaMK2rep2 and nCaMK2rep3 were immunoprecipitated using anti-myc antibody. Non-infected neurons were utilized as controls (Cnt). A clear interaction of nCaMK2rep3 sensor with endogenous PSD-95 protein was observed, dependent on the PSD95.FingR intrabody domain. From this point, nCaMK2rep3 was routinely used to infect cultured neurons and we refer to it as nCaMK2rep.

### CaMK2rep specifically reports CaMKII activity in cultured hippocampal neurons

Having established the sensitivity and specificity of CaMK2rep in a heterologous system, we evaluated its performance in a more physiological context. Cultured hippocampal neurons offer a native environment where endogenous CaMKII is highly expressed and dynamically regulated by synaptic activity. We therefore tested whether CaMK2rep could reliably report activity-dependent CaMKII activation in this neuronal setting.

To express CaMK2rep in cultured hippocampal neurons, we generated several lentiviral constructs under the control of the human synapsin promoter to ensure neuron-specific expression. Postsynaptic targeting was achieved by fusing a nanobody (PSD95.FingR), which selectively binds the postsynaptic scaffolding protein PSD95^45^, to the N-terminus of CaMK2rep. Figure 4A illustrates the four resulting constructs, designated nCaMK2rep-1 through –4, and their immunoblot analysis. Upon expression in neurons, rep1 and rep4 constructs showed low expression levels, whereas rep2 and rep3 showed markedly higher expression. Basal phosphorylation levels followed a similar trend, with rep2 and rep3 displaying stronger signals, likely due to their greater abundance. We next assessed the subcellular distribution of the constructs (Figure 4B). Constructs containing PSD95.FingR, such as rep4, exhibited a more punctate, dendrite-enriched localization compared to those lacking the nanobody, such as rep2, which were largely somatic. This pattern was initially evident from endogenous GFP fluorescence but became more pronounced upon immunostaining with the phospho-S603-specific antibody (D4B9I). Constructs lacking PSD95.FingR showed predominant somatic phosphorylation, while those containing it exhibited a punctate, dendritic distribution that overlapped with MAP2 staining. Notably, in non-infected neurons, the signal detected by the D4B9I antibody –which recognizes endogenous synapsin phosphorylated at phosphosite 3—was markedly weaker than in infected neurons, indicating that the majority of the pSer603 signal in infected cells originates from the nCaMK2rep sensor. To confirm interaction with endogenous PSD95, we performed immunoprecipitation experiments. As shown in Figure 4C, rep3 – bearing the PSD95.FingR domain-co-immunoprecipitated with PSD95, whereas rep2 did not. Based on these expression, localization, and interaction profiles, we selected rep3 for all subsequent neuronal experiments. This construct will hereafter be referred to as nCaMK2rep.

To validate the specificity of nCaMK2rep in cultured hippocampal neurons, we first examined its sensitivity to pharmacological inhibition of CaMKII. Application of the Ca^2+^/CaM-competitive inhibitor KN-93 significantly reduced basal phosphorylation of nCaMK2rep, while its inactive analog, KN-92, had no effect (Figure 5A). Treatment with Ruxolitinib –a recently described, potent, and selective CaMKII inhibitor^1,46^-further decreased nCaMK2rep phosphorylation, providing additional support for the reporter’s specificity toward CaMKII activity. Importantly, nCaMK2rep expression did not alter endogenous CaMKII levels in neuronal cultures. To further confirm specificity, we overexpressed CaMKII using a lentiviral vector, which led to a pronounced increase in nCaMK2rep phosphorylation. Similarly, expression of a photoactivatable CaMKII variant (paCaMKII) resulted in a more than twofold increase in reporter phosphorylation upon 2-minute exposure to 460 nm light (Figure 5B), whereas no change was observed in neurons expressing the light-insensitive control variant (paCaMKII-SD). These results confirm that nCaMK2rep reliably reports on CaMKII activation in neurons. Given the central role of CaMKII in synaptic plasticity –particularly downstream of NMDA receptor activation and calcium influx-we next tested whether nCaMK2rep could report activity induced by NMDA receptor stimulation. Neurons were exposed to three different NMDA receptor-dependent stimulation paradigms: (1) direct application of NMDA; (2) disinhibition via bicuculline combined with glycine and strychnine (BIC); and (3) application of the NMDA receptor positive allosteric modulators UBP684 and UBP714, which enhance receptor responses during spontaneous synaptic activity. In all conditions, we observed robust increases in nCaMK2rep phosphorylation, with the strongest responses elicited by UBP714 and the bicuculline/glycine/strychnine combination. Notably, these effects were absent in neurons expressing the non-phosphorylatable mutant reporter nCaMK2rep-S603A, further confirming the specificity of the phosphorylation signal. Together, these findings establish nCaMK2rep as a highly sensitive and specific reporter of CaMKII activity in hippocampal neurons, capable of detecting both basal and activity-dependent changes relevant to synaptic plasticity.

**Figure 5.**
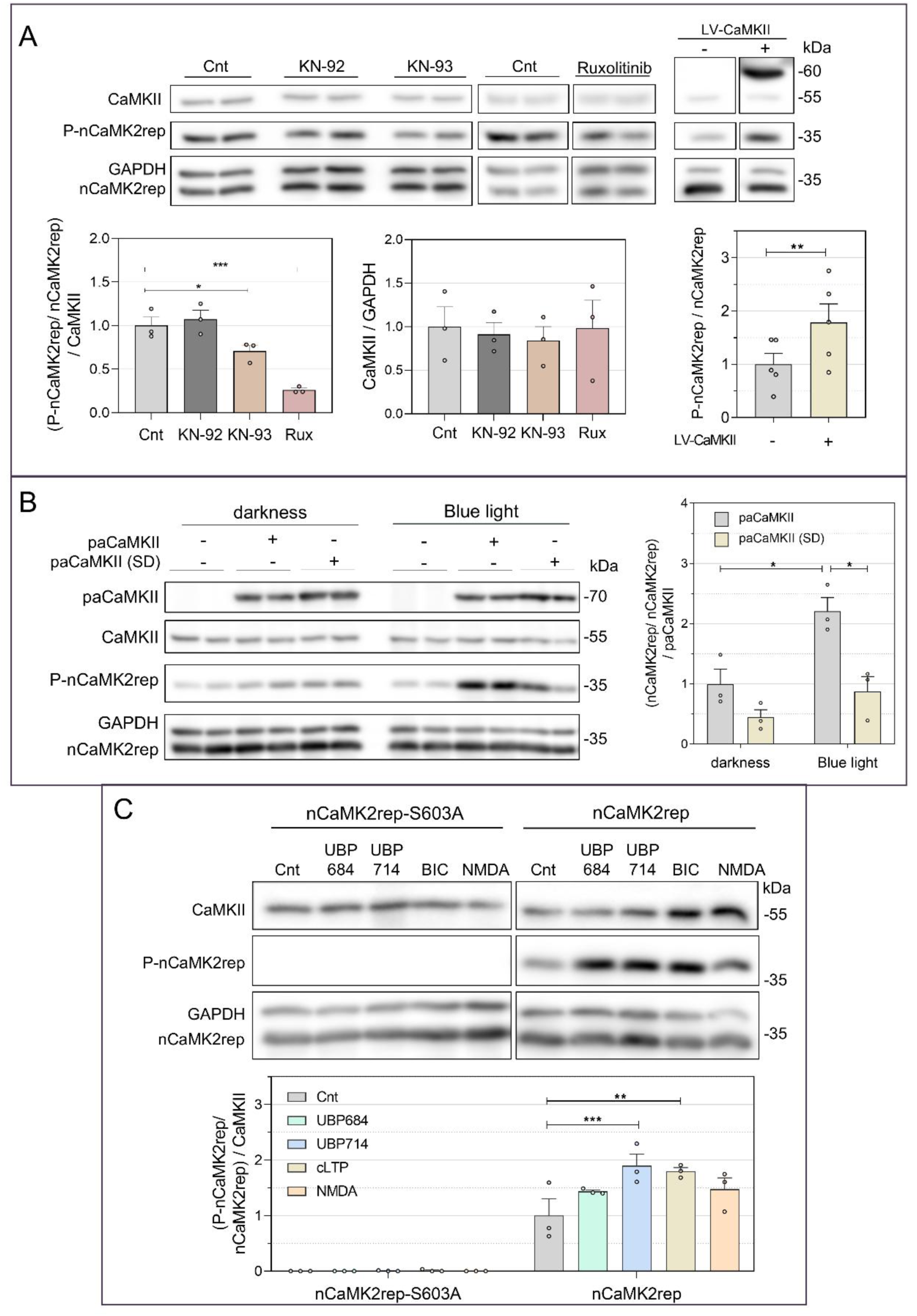
nCaMK2rep reports on basal and stimulated CaMKII activity in hippocampal neurons. **A.** Hippocampal neurons infected at DIV4 with nCaMK2rep were treated at DIV16 with 20 μM KN-93, its inactive analog KN-92, or 10 μM Ruxolitinib (10 minutes, 37°C). These treatments did not alter endogenous expression of CaMKII. In contrast, lentiviral expression of CaMKII from DIV4 increased nCaMK2rep phosphorylation levels (mean ± SEM, n=5). **B.** Hippocampal neurons expressing nCaMK2rep were infected at DIV7 with AAV-paCaMKII, and nCaMK2rep phosphorylation analyzed at DIV16 after either blue light stimulation (460 nm, 2 minutes) or maintained in darkness. paCaMKII (SD) – a light-insensitive form of paCaMKII – was used as a control (mean ± SEM, n=3). **C.** Hippocampal neurons expressing nCaMK2rep were stimulated for 5 minutes at DIV16 with one of the following treatments: 30 μM UBP684, 30 μM UBP714, 25 μM bicuculline (plus 200 μM glycine and 1 μM strychnine), or 20 μM NMDA, resulting in increased phosphorylation of nCaMK2rep. The nCaMK2rep-S603A mutant was used as a negative control for phosphorylation (mean ± SEM, n=3).

### Neurogranin, a CaM-binding protein, attenuates CaMKII activity

Neurogranin (Ng) –a postsynaptic protein we have previously studied extensively^47,48^-is highly abundant in the forebrain and plays a key role in synaptic plasticity by modulating CaM availability^49^. Ng binds CaM at low intracellular Ca^2+^ concentrations and releases it when Ca^2+^ rises; this interaction is blocked by PKC-mediated phosphorylation of Ng. As mentioned, these properties have led to two competing hypotheses: one suggests that Ng targets CaM to postsynaptic sites to facilitate rapid activation of effectors like CaMKII upon stimulation, while the other proposes that Ng buffers CaM, therefore limiting CaM-dependent signaling. Despite these differing views, Ng is clearly essential for normal brain function: mice lacking Ng show impaired LTP and deficits in hippocampus-dependent learning, highlighting its central role in synaptic signaling and cognition^50^. To shed light on these models, we examined how Ng expression affects CaMKII activity both in HeLa cells and in hippocampal neurons using our newly developed reporter.

We first examined CaMK2rep phosphorylation in basal conditions and following histamine (HA) stimulation, in HeLa cells co-expressing either wild-type Ng (Ng-wt) or Ng mutants with altered calmodulin (CaM) binding properties (Figure 6A). The non-phosphorylatable Ng-S36A mutant retains CaM binding, as this interaction is not regulated by PKC. In contrast, the Ng-I33Q and Ng-S36D mutants^51^ are unable to bind CaM. We found that only Ng-wt and Ng-S36A significantly reduced CaMK2rep phosphorylation –both in resting and HA-stimulated conditions-suggesting that Ng primarily limits CaMKII activation by sequestering CaM. To determine whether this mechanism also operates in neurons, we repeated the experiment in cultured hippocampal neurons expressing nCaMK2rep. Consistent with HeLa cell results, co-expression of Ng-wt or Ng-S36A significantly decreased reporter phosphorylation, whereas the CaM-binding-deficient mutants had no effect (Figure 6B). These findings support the notion that Ng-wt and Ng-S36A reduce CaMKII activation by restricting CaM availability. If this is the case, then reducing endogenous Ng levels should enhance CaMKII activity. To test this, we knocked down Ng using a specific AAV-shRNA (shNg), and found a 2.5-fold increase in nCaMK2rep phosphorylation (Figure 6C), consistent with an increased availability of free CaM.

**Figure 6.**
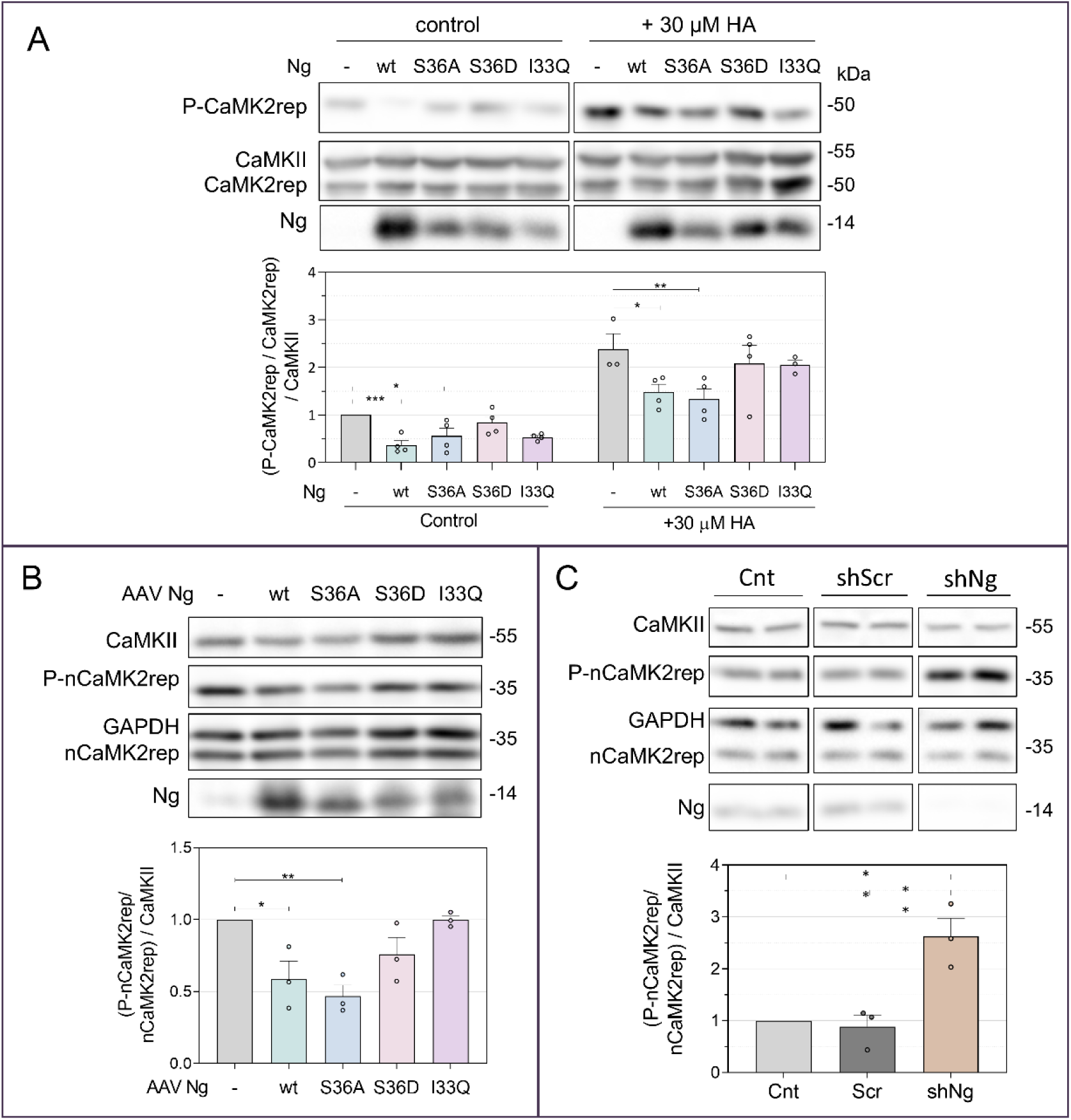
Ng attenuates CaMKII activity both in HeLa cells and hippocampal neurons. **A.** HeLa cells expressing CaMK2rep and CaMKII were further co-transfected with either wild-type Ng (Ng-wt) or the mutants Ng-S36A, Ng-S36D, and Ng-I33Q. 24 hours later, cells were stimulated with 30 µM HA for 2 minutes or left unstimulated. The histogram shows the levels of CaMK2rep referred to CaMKII expression and normalized to the control condition of no Ng expression (mean ± SEM, n=4). **B-C.** Hippocampal neurons expressing nCaMK2rep were infected at 7 DIV with AAVs to either express and mutants (**B**) or to silence endogenous Ng (**C**). AAVs expressing a scramble shRNA (Scr) was used as a control in the silencing experiments. Basal phosphorylation of nCaMK2rep was analyzed at DIV15-16 and normalized to the phosphorylation values obtained in the absence of Ng overexpression or silencing (mean ± SEM, n=3).

During the course of our study, a new genetically encoded CaMKII activity reporter, CaMKAR^1^, was published. This ratiometric sensor, based on the CaMKII autophosphorylation motif MHRQETVDCLK, follows the design principles of the ExRai-AKAR2 PKA biosensor^52^. According to the authors, CaMKAR offers enhanced dynamic range, faster kinetics, and greater sensitivity compared to previous FRET-based CaMKII biosensors^19,23^. To independently validate our findings on Ng-mediated modulation of CaMKII activity, we used CaMKAR in live-cell imaging experiments, both in HeLa cells and cultured hippocampal neurons. In HeLa cells, HA stimulation induced a rapid increase in CaMKAR 470/405 fluorescence ratio (ΔF/F₀), followed by a gradual decay (Figure 7A). This response was markedly reduced in cells co-expressing Ng-wt or the non-phosphorylatable, CaM-binding mutant Ng-S36A. In contrast, the CaM-binding-deficient mutants Ng-S36D and Ng-I33Q had no effect. As a negative control, we used CaMKAR-T6A, a non-phosphorylatable variant in which the critical threonine residue is replaced by alanine. A similar pattern was observed when analyzing CaMKAR activation in hippocampal neurons stimulated with glutamate (Figure 7B): Ng-wt and Ng-S36A attenuated CaMKAR responses, while Ng-S36D and Ng-I33Q did not. Interestingly, unlike with nCaMK2rep, Ng knockdown (shNg) did not enhance CaMKAR responses in neurons. This discrepancy may reflect differences in experimental paradigms –glutamate-induced responses for CaMKAR versus basal/spontaneous activity measured by nCaMK2rep. Each trace in Figure 7 represents the averaged response from multiple individually analyzed neurons, normalized to each cell’s baseline (ΔF/F₀ of the ex470/405 ratio). Because Ng and its mutants were not fluorescently tagged, we performed post hoc immunostaining to verify co-expression with nCaMK2rep. We found that over 90% of CaMKAR-positive neurons were also positive for Ng in all experimental groups (Ng-wt: 94.29 ± 4.16%; Ng-S36A: 95.57 ± 2.93%; Ng-S36D: 90.17 ± 5.16%; Ng-I33Q: 92.53 ± 2.63%; mean ± SEM, n = 6–7).

**Figure 7.**
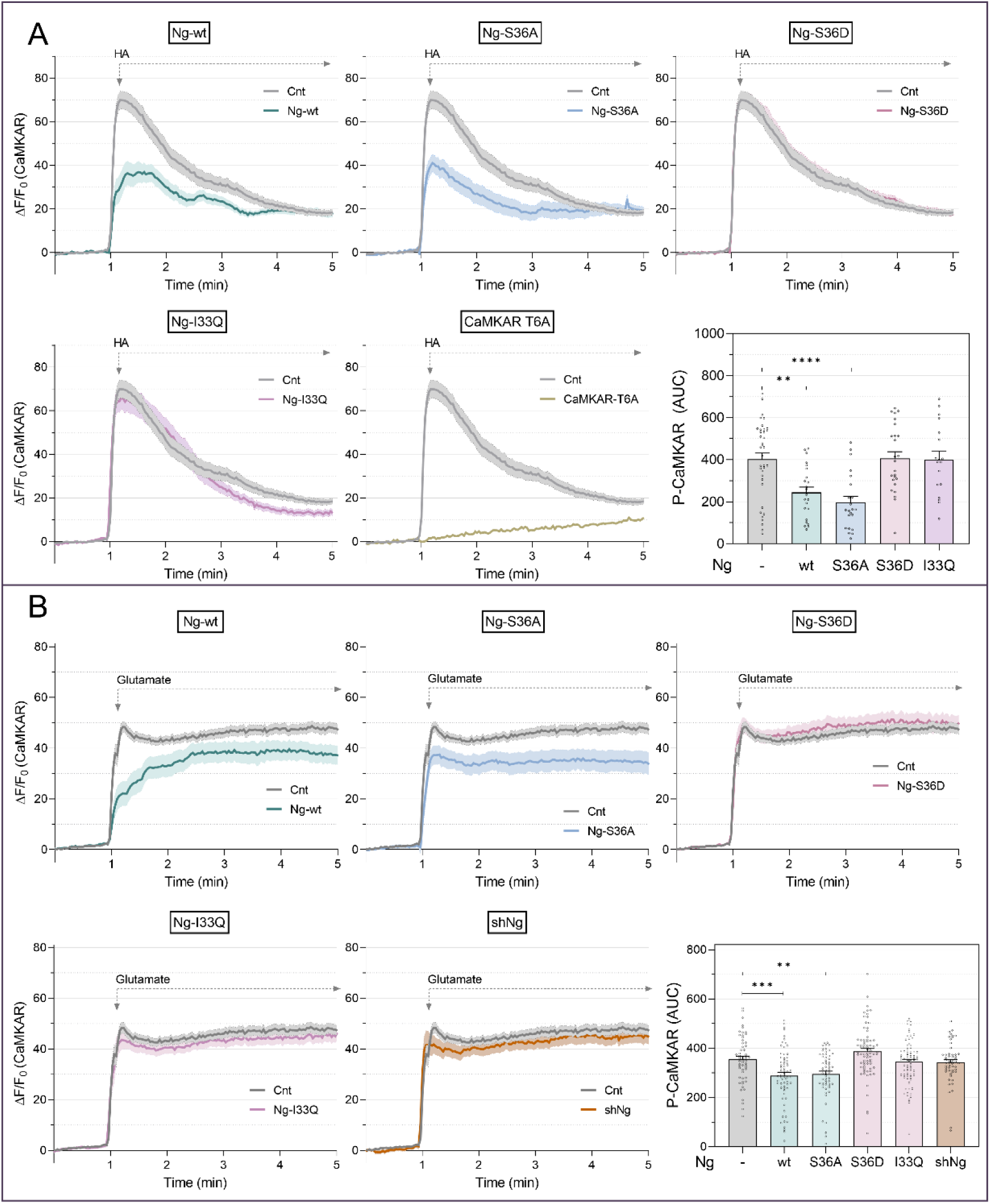
CaMKAR, a novel live-cell reporter of CaMKII activity, confirms Ng attenuation of CaMKII activity. **A-B**. CaMKAR phosphorylation was analyzed in HeLa cells (**A**) and in DIV16 hippocampal neurons (**B**) expressing (n)CaMKAR only (Cnt) or (n)CaMKAR along with Ng-wt or the mutants Ng-S36A, Ng-S36D, and Ng-I33Q. In HeLa cells, CaMKII was additionally expressed. In neurons, nCaMKAR phosphorylation was also assessed following Ng silencing from DIV10. The CaMKAR-T6A sensor was used as a negative control for phosphorylation. CaMKAR phosphorylation signal is expressed as the increase of the fluorescence ratio obtained upon excitation at 470 nm and 405 nm ΔF/F_0_ over the baseline F_0_, which is the mean of the ratios obtained during the first 60 seconds before the addition of 30 µM HA in HeLa cells (**A**) or 100 µM glutamate in hippocampal neurons (**B**). (n)CaMKAR phosphorylation was assessed as the area under the curve during the 2-minute period following treatment. **A**: Mean ± SEM, n = 20–37 cells. **B**: Mean ± SEM, n = 59–73 cells.

## Discussion

CaMKII is a central signaling molecule in neurons, translating calcium influx into long-lasting changes in synaptic strength^2^. Understanding the spatial and temporal dynamics of its activation is essential for unraveling the molecular basis of synaptic plasticity and its involvement in cardiovascular and neurological disorders. Here, we introduce and validate CaMK2rep, a genetically encoded phosphorylation reporter specifically designed to monitor CaMKII activity with high sensitivity and specificity. CaMK2rep incorporates two synapsin1a-derived CaMKII phospho-epitopes recognized by a commercially available antibody. This non-FRET, phosphorylation-dependent reporter offers a robust and versatile tool for detecting CaMKII activation under basal and stimulated conditions. The neuronal version, nCaMK2rep, allows targeted analysis within excitatory postsynaptic compartments and reveals endogenous CaMKII activity patterns in neurons. Using this tool, we investigated the role of neurogranin (Ng), a highly abundant CaM-binding protein enriched in dendritic spines of forebrain excitatory neurons^49^, in modulating CaMKII signaling. Our findings consistently support a model in which Ng functions as a CaM buffer, limiting CaMKII activation by reducing the availability of free CaM under both resting and stimulated conditions. These results were independently validated using CaMKAR, a ratiometric, non-FRET-based live-cell reporter, reinforcing the robustness of our observations and demonstrating the utility of CaMK2rep in probing CaMKII signaling in neurons.

Compared to existing CaMKII activity sensors, CaMK2rep offers several advantages. Unlike FRET-based reporters, which require specialized imaging equipment and often suffer from limited dynamic range, CaMK2rep relies on phosphorylation-dependent antibody detection, making it compatible with standard immunoblotting and immunofluorescence approaches that offer high sensitivity and signal amplification. This compatibility enables multiplexed biochemical and morphological analyses across multiple samples or conditions. When combined with temporally controlled stimulation protocols, CaMK2rep also retains sufficient temporal resolution to resolve activity dynamics. A potential concern in neurons is that phosphorylated synapsin at phosphosite 3 might contribute to the detected signal and complicate interpretation in immunofluorescence analysis. This is not an issue for immunoblotting, where the signal from synapsin and CaMK2rep are clearly separated. As shown in Figure 4B, endogenous synapsin phosphorylation at phosphosite 3 accounts for only a small fraction of the overall signal observed upon nCaMK2rep expression in cultured hippocampal neurons. To eliminate any residual contribution from endogenous synapsin in immunofluorescence, proximity ligation assays (PLA) can be employed using antibodies against phospho-synapsin (D4B9I) and the Myc epitope. Since CaMK2rep contains three consecutive Myc tags at its C-terminus, this strategy enables selective detection of reporter-specific phosphorylation events.

Validation experiments confirmed the high specificity of CaMK2rep for CaMKII. In HeLa cells, basal phosphorylation of the reporter required co-expression of CaMKIIα and was strongly enhanced by calcium-elevating stimuli. Reporter activation was blocked by pharmacological inhibitors KN-93 and Ruxolitinib, and was absent when unrelated kinases such as PKA or PKC were expressed. Interestingly, KN-93 was more effective in HeLa cells than in neurons, likely reflecting a higher fraction of T286-autophosphorylated, autonomously active CaMKII in neurons –a form resistant to KN-93 inhibition^53^. This notion is further supported by the stronger inhibition observed in neurons with Ruxolitinib. CaMK2rep also responded to activation of a photoactivatable CaMKII construct, further supporting its selectivity. In hippocampal neurons, the reporter retained sensitivity and spatial resolution, particularly when targeted to postsynaptic compartments via PSD95.FingR. The neuronal variant, nCaMK2rep, detected both basal and NMDA receptor-induced CaMKII activity and remained responsive under pharmacological and genetic manipulations. Importantly, CaMK2rep expression showed no signs of toxicity: HeLa cells maintained its typical morphology and proliferated normally, and hippocampal neurons exhibited no deficits in development or dendritic arborization. These features position CaMK2rep as a reliable tool for precise monitoring of endogenous CaMKII signaling.

Among alternative approaches that could compete with CaMK2rep, a widely used proxy for CaMKII activity is the detection of T286 autophosphorylation using phospho-specific antibodies –a method technically similar to ours. However, this approach detects only the Ca²⁺-independent, autonomously active form of CaMKII, which requires prior activation of adjacent subunits by Ca²⁺/CaM and retains just ∼20–40% of maximal kinase activity^54,55^. It does not capture activity from Ca^2+^/CaM-bound subunits that have not undergone T286 phosphorylation, nor from GluN2B-bound CaMKII, which is independent of both Ca^2+^/CaM and T286 autophosphorylation^16^. Additionally, in situ immunofluorescence detection may be limited, as pT286-CaMKII localizes to the postsynaptic density, a dense protein matrix that can hinder antibody access. Other approaches involve fluorescent protein-based biosensors, which are powerful tools for live-cell imaging of signaling dynamics. However, they often suffer from limited sensitivity and dynamic range. CaMK2rep addresses these limitations by incorporating two identical synapsin phosphosite 3 sequences flanked by a native sequence context, greatly enhancing its sensitivity. Indeed, dual-site constructs produced substantially stronger signals than single-site variants, as evidence in this study. Although it remains unclear which of the two sites is preferentially phosphorylated or more accessible to antibody detection –or how this varies across experimental conditions-CaMK2rep exhibited an almost linear response to both CaMKII expression levels and stimulation intensity, underscoring its versatility and utility. Its compatibility with immunoblotting enables population-level quantification across multiple conditions (e.g., drug screening), while its use in immunofluorescence provides spatially resolved readouts at the cellular and subcellular level. These features make CaMK2rep a practical and scalable reporter for high-content or multiplex analysis in complex signaling studies.

We next used CaMK2rep to investigate the functional role of Ng^56^ in CaM signaling. Previous models propose that Ng either acts as a CaM-targeting scaffold to promote rapid CaMKII activation upon synaptic stimulation or as a CaM buffer that limits CaM availability under low-Ca^2+^ conditions^57,58^. Our data support the latter: expression of wild-type Ng or the non-phosphorylatable CaM-binding mutant S36A reduced CaMKII activity, while CaM-binding-deficient mutants S36D and I33Q had no effect. Conversely, knockdown of endogenous Ng in neurons led to increased nCaMK2rep signal, supporting a negative regulatory role. This pattern was independently confirmed using CaMKAR. Together, these findings provide direct evidence that Ng acts primarily as a CaM-sequestering protein, attenuating CaMKII activation in resting or weakly stimulated neurons. Nonetheless, alternative mechanisms cannot be entirely excluded. For example, a computational model by Li et al.^59^ suggests that Ng’s role may depend on stimulation frequency: at intermediate frequencies, it acts as a CaM buffer, whereas at high frequencies, it may inhibit calcineurin and enhance CaMKII autophosphorylation.

In summary, we present CaMK2rep as a robust, sensitive, and specific tool for monitoring CaMKII activity across diverse cellular contexts. Validated in both heterologous systems and neurons, and successfully applied to a longstanding question in synaptic signaling, CaMK2rep proves valuable not only as a biochemical readout but also as a mechanistic probe of CaM-dependent regulation. Importantly, our findings highlight the complementary strengths of CaMK2rep and CaMKAR^1^. CaMK2rep, which relies on antibody-based detection, enables sensitive biochemical and detailed morphological analyses, while CaMKAR permits real-time monitoring of CaMKII activity in live cells. This synergy favors conducting preliminary population analyses with CaMK2rep that inform subsequent live-cell studies with CaMKAR, thereby helping to delve into the mechanistic understanding of CaMKII activity across various contexts. Together, these reporters provided complementary insights into Ng function in synaptic plasticity. Our findings support a model in which Ng primarily acts as a CaM-sequestering protein that limits basal CaMKII activation. More broadly, this study illustrates the power of genetically encoded sensors in resolving persistent controversies in signal transduction. CaMK2rep adds a valuable new tool to the repertoire for studying CaMKII signaling in physiological and pathological contexts, including synaptic plasticity.

## Author Statements & Declarations

### Availability of data and materials

The plasmids generated in this study are publicly available at Addgene. CaMK2rep (Addgene # 239622), CaMK2rep-S603A (Addgene # 239623), nCaMK2rep (Addgene # 239624), nCaMK2rep-S603A (Addgene # 239625).

### Competing interests

Authors Elena Martínez-Blanco (E.M-B.), Raquel de Andrés (R.A.), Lucía Baratas-Álvarez (L.B-A.) and F. Javier Díez-Guerra (F.J.D-G.) declare they have no commercial interests.

### Funding

The author Elena Martínez-Blanco (E.M-B.) received research support from the “Comunidad de Madrid”, in the form of contract PEJD-2018/PRE/BDM-8491. The author Raquel de Andrés (R.A.) received research support from the “Comunidad de Madrid”, a regional public institution, in the form of contracts PEJ-2017-AI/BMD-6017 and PEJD-2019-PRE/BMD-14947. The author Lucía Baratas-Álvarez (L.B-A.) received research support from the “Comunidad de Madrid”, in the form of contract TL-CM PJ 2019 TL-BMD-13238. E.M-B., R.A. and L.B-A. were supported by the “Fundación Severo Ochoa”, a private organization ascribed to the Center for Molecular Biology (CSIC-UAM). The present work was initially supported by the Spanish Ministry of Science, Innovation and Universities (grant RTI2018-098712-B-I00).

### Author Contributions

Elena Martínez-Blanco (E.M-B.) and Raquel de Andrés (R.A.) carried out the experiments, analyzed the data and participated in discussions. Lucía Baratas-Álvarez (L.B-A.) provided technical support. F. Javier Díez-Guerra (F.J.D-G.) conceived the study, contributed to data analysis and wrote the manuscript. All authors have read and approved the final version of the manuscript.

### Ethics approval

All procedures were carried out in accordance with the Spanish Royal Decree 1201/2005 for the protection of animals used in scientific research and the European Union Directive 2010/63/EU regarding the protection of animals used for scientific purposes. The procedures were approved by the local institute and regional local ethics committees (“Comunidad de Madrid”, Ref. PROEX 106/19).

## Acknowledgements

We thank the Advanced Light Microscopy Core Facility (SMOA) of the “Centro de Biología Molecular” (CBM) for assistance with the imaging studies. We thank “Fundación Ramón Areces” for providing institutional support to CBM.

## Figure Legends

**Supplementary Figure 1.**
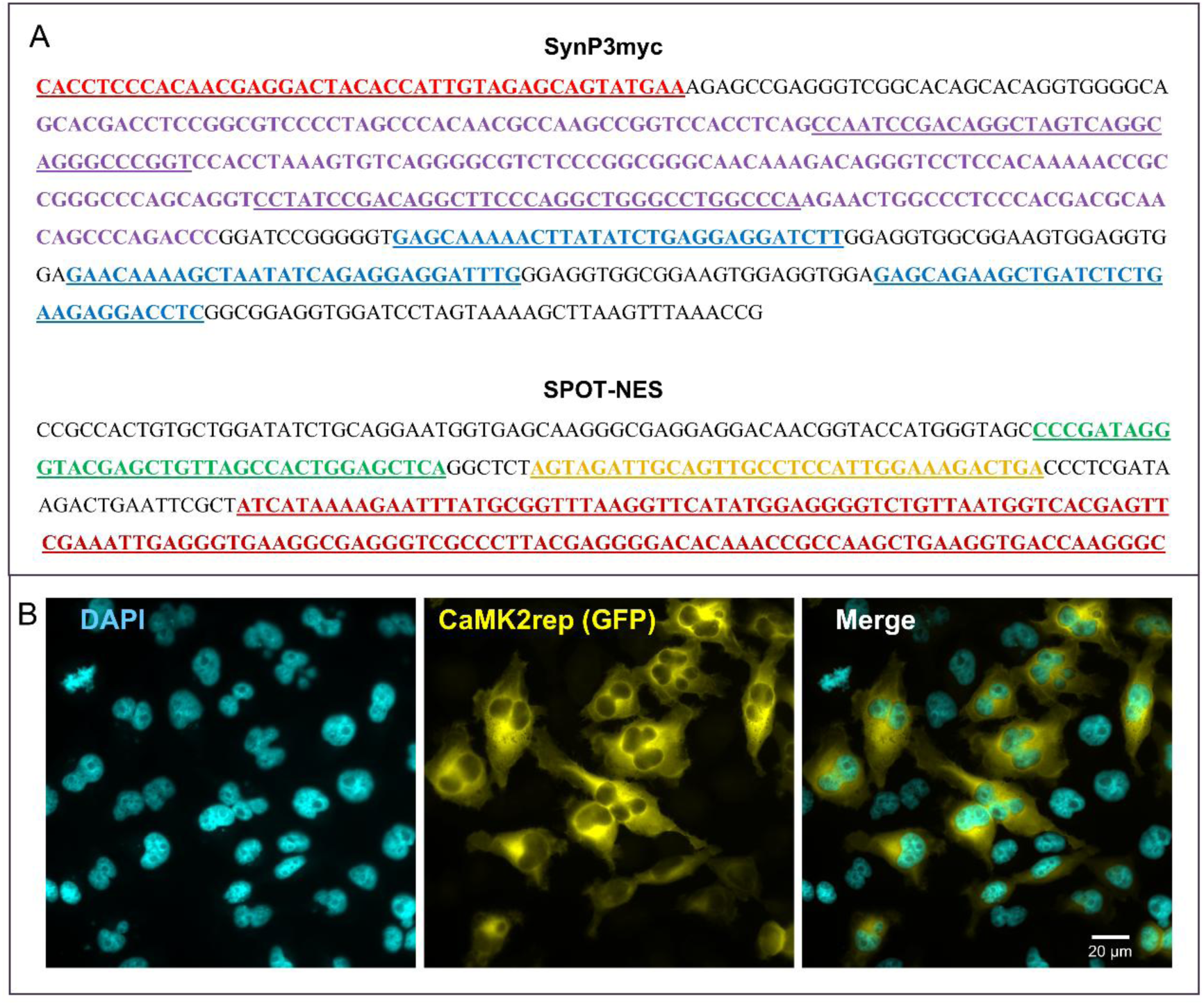
Design of CaMK2rep, a CaMKII activity reporter. **A.** SynP3myc is a synthetic DNA fragment that included 43 bp from mCherry C-terminal (red), 237 bp from a cDNA coding for amino acid sequence 543-620 from rat synapsin-1a (purple) with two copies of CaMKII-specific phospho-site 3 (underlined) and three consecutive copies of the myc tag (blue). Spot-NES sequence contains a Spot-tag (green), a nuclear export sequence (NES, orange) and 135 bp of the C-terminal sequence of mCherry (red). **B.** Immunofluorescence of HeLa cells transfected with CaMK2rep, with nuclei stained using DAPI. Images were acquired in the DAPI (ex378/52; em432/36) and GFP (ex474/27; em515/30) channels, with a 63X NA 1.4 objective in a Zeiss Axiovert 200M fluorescence microscope.

**Supplementary Table 1.** Plasmids developed and used in this study.

| Plasmids | Backbone | Insert | promoter | expression | tag |
| --- | --- | --- | --- | --- | --- |
| CaMK2rep | pcDNA3.1 | SpotNES-sGFP2-SynP3-3xmyc | CMV | Mammalian | sGFP2, myc |
| CaMKIIα | pcDNA3<br>pLOX | CaMKIIα-3xmyc | CMV<br>synapsin | Mammalian<br>lentiviral | myc |
| nCaMK2rep1 | pLOX | NES-SynP3-3xmyc | synapsin | lentiviral | myc |
| nCaMK2rep2 | pLOX | NES-sGFP2-SynP3-3xmyc | synapsin | lentiviral | sGFP2 |
| nCaMK2rep3 | pLOX | PSD95.FingR-NES-SynP3-3xmyc | synapsin | lentiviral | myc |
| nCaMK2rep4 | pLOX | PSD95.FingR-NES-sGFP2-SynP3-3xmyc | synapsin | lentiviral | sGFP2, myc |
| tdTomato-P2A-jGCaMP8s | pcDNA3 | tdTomato-P2A-jGCaMP8s | CMV | Mammalian | tdTomato |
| PKA-C-α-YFP* | pEYFP-N2 | | CMV | Mammalian | YFP |
| PKCγ-YFP* | pEYFP-N2 | | CMV | Mammalian | YFP |
| Ng | pcDNA3<br>pAAV | Ng | CMV<br>CaMKIIα0.4 | Mammalian<br>AAV2 | |
| Ng-S36A | pcDNA3<br>pAAV | Ng-S36A | CMV<br>CaMKIIα0.4 | Mammalian<br>AAV2 | |
| Ng-S36D | pcDNA3<br>pAAV | Ng-S36D | CMV<br>CaMKIIα0.4 | Mammalian<br>AAV2 | |
| Ng-I33Q | pcDNA3<br>pAAV | Ng-I33Q | CMV<br>CaMKIIα0.4 | Mammalian<br>AAV2 | |
| pAAV-shNg-mRuby2 | pAAV | shRNA targeting Ng | H1 | AAV2 | mRuby2 |
| pFΔ6** | | AAV helper | | AAV2 | |
| pRV1** |  | cap |  | AAV2 |  |
| pH21** |  | rep-cap |  | AAV2 |  |

\* kindly donated by Dr Mark Dell'Aqua (Department of Pharmacology, University of Colorado, Denver, USA)
\*\* kindly donated by Dr Hilmar Bading (Department of Neurobiology, Interdisciplinary Center for Neurosciences (IZN), Heidelberg, Germany)

**Supplementary Table 2.** Plasmids obtained from Addgene.

| Plasmids | Addgene | Backbone | Insert | promoter | expression | tag |
| --- | --- | --- | --- | --- | --- | --- |
| CMV-tdTomato-P2A-paCaMKII | #165431 |  | tdTomato-P2A-paCaMKII | CMV | Mammalian | tdTomato |
| CMV-tdTomato-P2A-paCaMKII K42M | #165433 |  | tdTomato-P2A-paCaMKII-K42M | CMV | Mammalian | tdTomato |
| CaMKAR* | #205315 | pcDNA3 | CaMKAR | CMV | Mammalian |  |
| pAAV-CaMP0.4-FHS-paCaMKII-WPRE3 | #165429 | pAAV | FHS-paCaMKII | CaMKIIα0.4 | AAV2 | FHS (Flag, HisX6, Strep) |
| pAAV-CaMP0.4-FHS-paCaMKII(SD)-WPRE3 | #165430 | pAAV | FHS-paCaMKII(SD) | CaMKIIα0.4 | AAV2 | FHS (Flag, HisX6, Strep) |
| pAAV-shNg-mRuby2 | Modified from #92155 | pAAV | shRNA targeting Ng | H1 | AAV2 | mRuby2 |
| pAAV-U6_CaMKIIa-mCherry | #181875 | pAAV | control shRNA | U6 | AAV2 | mCherry |
| pCMVRδ8.74 | #22036 | | gag pol tat rev | | lentiviral | |
| pMD2.G | #12259 |  | VSV-G envelope |  | lentiviral |  |

\* kindly donated by Dr. Jonathan M. Granger (Johns Hopkins University School of Medicine, Baltimore, USA)

**Supplementary Table 3.**
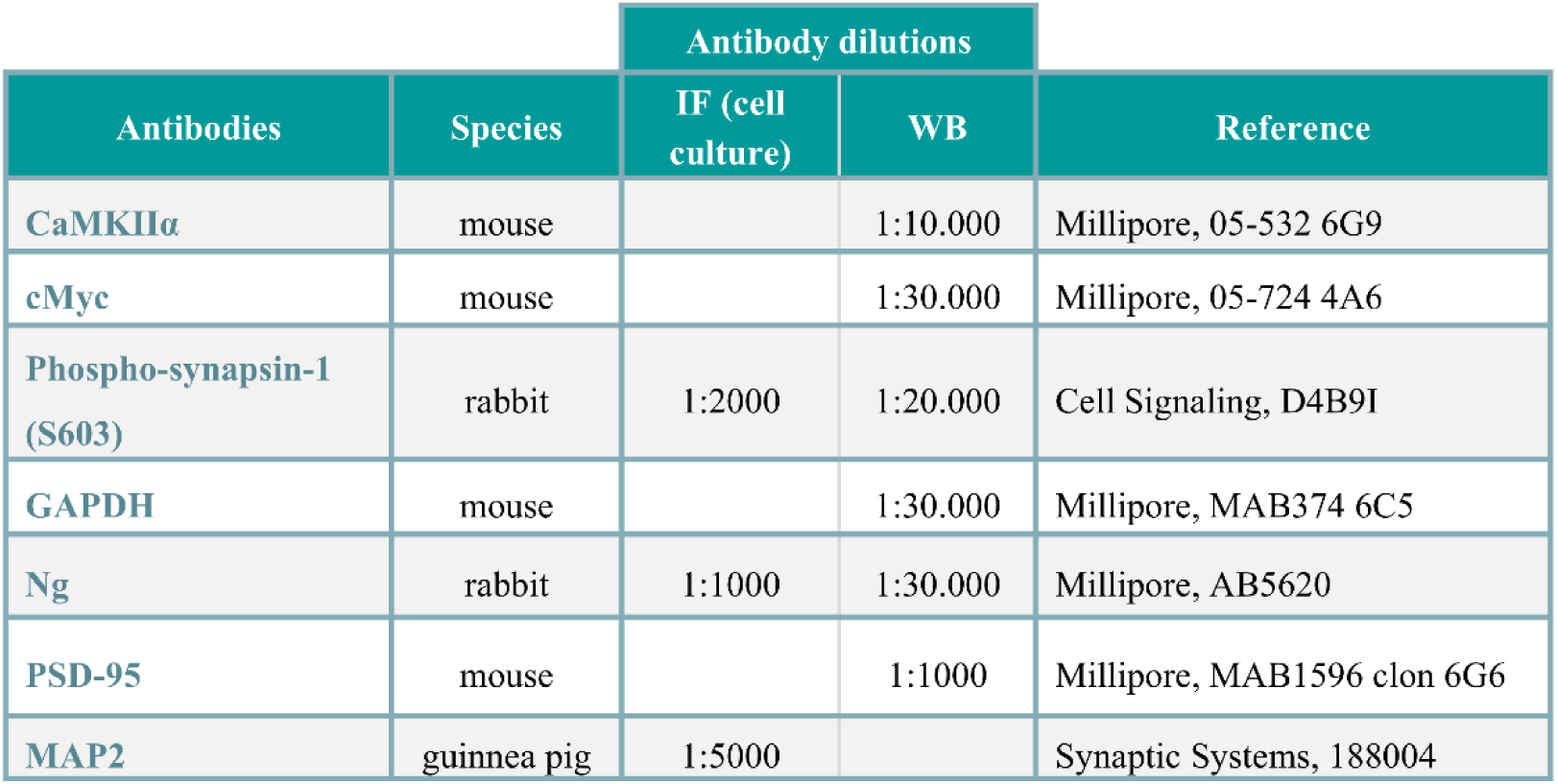
Antibodies used in this study for WB and IFs.

| Antibodies | Species | Antibody dilutions |  | Reference |
| --- | --- | --- | --- | --- |
|  |  | IF (cell culture) | WB |  |
| CaMKIIα | mouse | | 1:10.000 | Millipore, 05-532 6G9 |
| cMyc | mouse |  | 1:30.000 | Millipore, 05-724 4A6 |
| Phospho-synapsin-1 (S603) | rabbit | 1:2000 | 1:20.000 | Cell Signaling, D4B9I |
| GAPDH | mouse |  | 1:30.000 | Millipore, MAB374 6C5 |
| Ng | rabbit | 1:1000 | 1:30.000 | Millipore, AB5620 |
| PSD-95 | mouse |  | 1:1000 | Millipore, MAB1596 clon 6G6 |
| MAP2 | guinea pig | 1:5000 |  | Synaptic Systems, 188004 |

## Notes

### Competing Interest Statement

The authors have declared no competing interest.

